# Natural variations within the glycan shield of SARS-CoV-2 impact viral spike dynamics

**DOI:** 10.1101/2022.08.17.504157

**Authors:** Maddy L. Newby, Carl A. Fogarty, Joel D. Allen, John Butler, Elisa Fadda, Max Crispin

**Affiliations:** School of Biological Sciences, University of Southampton, Southampton, UK; Department of Chemistry and Hamilton Institute, Maynooth University, Maynooth, Kildare, Ireland

**Keywords:** SARS-CoV-2, glycosylation, variant of concern, mass spectrometry, molecular dynamics

## Abstract

The emergence of SARS-CoV-2 variants alters the efficacy of existing immunity, whether arisen naturally or through vaccination. Understanding the structure of the viral spike assists in determining the impact of mutations on the antigenic surface. One class of mutation impacts glycosylation attachment sites, which have the capacity to influence the antigenic structure beyond the immediate site of attachment. Here, we compare the glycosylation of a recombinant viral spike mimetic of the P.1 (Gamma) strain, which exhibits two additional N-linked glycan sites compared to the equivalent mimetic of the Wuhan strain. We determine the site-specific glycosylation of these variants and investigate the impact of these glycans by molecular dynamics. The N188 site is shown to exhibit very limited glycan maturation, consistent with limited enzyme accessibility. Structural modeling and molecular dynamics reveal that N188 is located within a cavity by the receptor binding domain, which influences the dynamics of these attachment domains. These observations suggest a mechanism whereby mutations affecting viral glycosylation sites have a structural impact across the antigenic surface.

## Introduction

The emergence and rapid spread of severe acute respiratory syndrome coronavirus 2 (SARS-CoV-2) has led to a protracted global health emergency, with over 6 million deaths worldwide as of June 2022. The development and deployment of several vaccines using derivatives of the trimeric spike (S) glycoprotein based on the Wuhan-hu-1 lineage has subdued symptom severity and mortality through the elicitation of neutralizing antibodies (NAbs) and T-cell mediated protection [1–7]. Mechanisms of immune escape by SARS-CoV-2 are dominated by the accumulation of mutations within the S glycoprotein. Alterations to the antigenic surface of the S protein, the predominant target of NAbs, enables continuous circulation of the virus throughout the global population. Neutralization resistance to sera from both convalescent and vaccinated individuals has resulted in particular SARS-CoV-2 variants being termed variants of concern (VOC) [8–10]. These VOCs have prolonged the pandemic and has resulted in several rebounds in case numbers and deaths despite widespread vaccine uptake, and extensive transmission through the population. As such, understanding how pre-existing immunity is being impacted by these VOCs is important for preparing for the ongoing evolution of the SARS-CoV-2 virus.

The SARS-CoV-2 S protein is a trimeric class I fusion protein that decorates the surface of the virus and mediates viral infection [11,12]. Each protomer is comprised of an S1 and S2 subunit, which both contain functionally distinct protein domains [12]. The S1 subunit facilitates receptor binding and encompasses the N-terminal domain (NTD) and receptor binding domain (RBD). Viral fusion and cell entry are organized by the S2 subunit, which includes the fusion peptide (FP), heptad repeat domains (HR1 and HR2), central helix (CH), connector domain (CD), and transmembrane domain (TM). Several studies from sera of infected individuals have revealed that the spike glycoprotein, specifically the RBD, contains epitopes for neutralizing antibodies, and is therefore the focus of vaccine design efforts [13– 22]. SARS-CoV-2 has been subjected to immunological pressure, creating selective pressure for structural mechanisms that thwart vaccine- or infection-mediated viral sterilization, thereby enhancing immune evasion. The principle mechanism of immune evasion is the introduction of amino acid substitutions and indels within the S glycoprotein that hinder NAb recognition [23,24]. This may subsequently introduce or remove sequence-encoded asparagine (N) -linked glycan sites. Ostensibly immunologically self N-linked glycans can serve to efficiently shield the underlying immunogenic viral protein surface, hence any changes in the glycan shield resulting from emergent lineages warrant investigation.

Lineages B.1.351 (Beta) and P.1 (Gamma) became prominent within the global population from February 2021 and B.1.617.2 (Delta) in May 2021 [25]. These SARS-CoV-2 VOCs present increased transmissibility and pathogenicity, predominantly resulting from mutations arising within S1 and the receptor binding domain (RBD) [26–29]. The mutational landscape of these VOCs are vast, yet a distinct set of advantageous mutations have prevailed throughout the evolution of distinct lineages (**Figure 1**) [23,30]. Contributing to enhanced infectivity and ACE2 binding, the early emerging VOCs, B.1.351 and P.1, accrued RBD mutations K417N/T, E484K and N501Y which have cumulatively demonstrated decreased neutralization sensitivity and increased infectivity [29,31,32]. The introduction of mutations within this region can facilitate vaccine escape as a large portion of NAb responses generated against SARS-CoV-2 are directed towards the RBD, and most others directed towards other epitopes within S1 [33].

**Figure 1:**
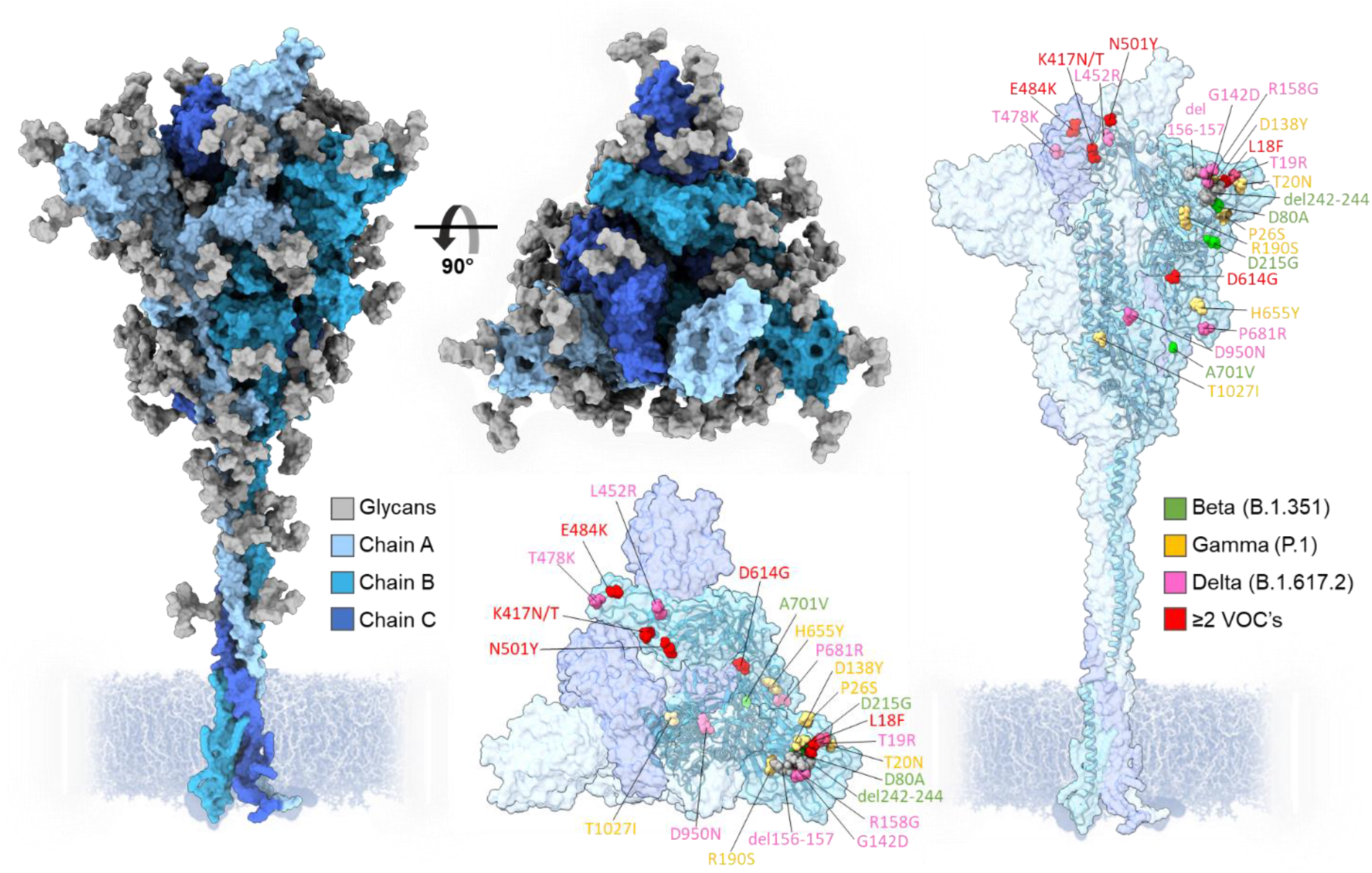
Three-dimensional representation of the SARS-CoV-2 S protein, showing regions of the spike that differ from the Wuhan strain in VOCs Beta, Gamma and Delta. Each protomer is colored in different shades of blue and N-linked glycans colored in gray. Amino acid mutations are color coordinated into lime green, Beta; yellow, Gamma; pink, Delta; red, ≥2 VOCs. Model of the Wuhan-hu-1 S glycan shield by Zuzic et al. 2022 [34].

Whilst the impact of the additional mutations present in the S protein variants has been well studied, there are potential for changes to occur that impact protein dynamics[35–38]. Previous studies have demonstrated that the SARS-CoV-2 S protein gene encodes for the attachment of several asparagine (N) linked glycosylation sites, with glycans accounting for approximately a third of the mass of fully mature S. [39–42] As such, it is important to monitor the changes in glycosylation that may occur in the S protein variants. Localized changes in the antigenic surface of the S glycoprotein can induce changes in the surrounding glycan architecture to potentiate immune evasion and infectivity[43].

N-linked glycosylation is an enzyme-catalyzed co-translational modification whereby a glycan precursor is covalently attached to asparagine residues within N-X-S/T sequons (X ≠ P). The early mammalian glycan processing pathway follows a linear trajectory whereby a Glc_3_Man_9_GlcNAc_2_-Asn is subjected to ER- and *cis-*Golgi-resident glucosidases and mannosidases, yielding aMan_5_GlcNAc_2_-Asn (Man5) glycan. The transfer of a β1,2-linked GlcNAc to the terminal D1 mannose residue, generates an intermediate glycan, GlcNAc_1_Man_5_GlcNAc_2_-Asn which facilitates the diversification of N-liked glycan processing and maturation through the *medial*- and *trans*-Golgi. Subsequent processing by this highly regulated and efficient translational modification pathway generates a heterogenous plethora of glycoforms that decorate most mammalian glycoproteins.

SARS-CoV-2, like many other viruses, hijack this process to shield its immunogenic protein surface from the host immune system [44–49]. Large abundances of immaturely processed glycans are rarely observed on cell surface mammalian glycoproteins [50], but such glycan signatures are present at sites of steric influence within SARS-CoV-2 S protein, such as N234 at the trimer interface [51,52]. Glycan processing can be sensitive to local tertiary and quaternary protein and glycan architecture. An abundance of under processed oligomannose-type glycans can arise if access to the glycan substrate by ER and *cis*-Golgi α-mannosidase is impeded, either through protein-glycan or glycan-glycan clashes [53]. This glycan processing prohibition signature is a reliable marker of native-like protein folding [54,55]. Most sites across the SARS-CoV-2 S protein, however, do present a large abundance of fully processed complex-type glycans [39,41,56,57].

Interestingly, the presence of T20N and R190S in the P.1 lineage introduces two potential N-linked glycosylation sites (PNGS) within S1, at positions N20 and N188. However, the accruement of N-linked glycan sites in the P.1 lineage is not unique to SARS-CoV-2 S glycoprotein evolution. This phenomenon has been previously demonstrated through the episodic emergence of ∼5 N-linked glycan sites in Influenza A H3 lineages over a period of 50 years, before reaching a hypothesized “glycan limit” [58]. Further, extended exposure of HIV-1 to the host immune system encourages the shift and overall accumulation of up to 5 PNGS in gp120 and outer domain regions, indicating that there must be adaptation to large survival pressures during chronic infection [59]. However, the location of the new glycan sites found in SARS-CoV-2 P.1 does not seem to occlude clustered sites of vulnerability, such as those targeted by NTD-targeting NAbs [60]. Additionally, the B.1.617.2 strain accumulates a T19R mutation, which abolishes the N17 PNGS. Whether this is an advantageous evolutionary mutation remains unknown.

Here, we have generated recombinant versions of the S proteins of B.1.351, P.1 and B.1.617.2 variants of SARS-CoV-2 to facilitate analysis of the glycan shield by mass spectrometry and interrogate the possible mechanistic role of the novel N188 glycan in the P.1 variant. To ensure analogous glycosylation between the recombinant proteins and viral-derived material, we introduced identical stabilizing proline mutations as previously applied to the Wuhan-hu-1 strain, termed HexaPro modifications (**Figure 2)** [61].

**Figure 2.**
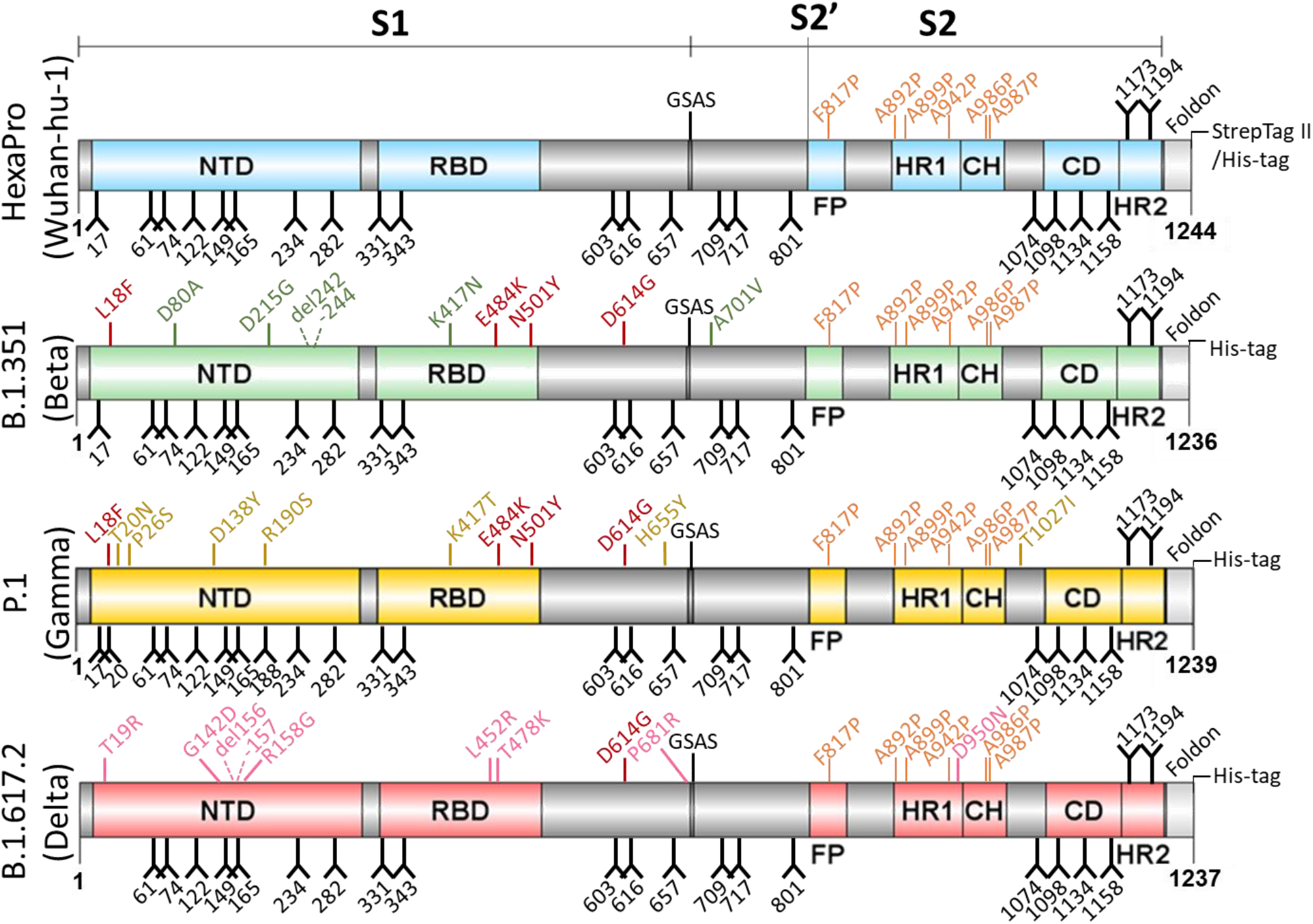
Schematic of soluble recombinant SARS-CoV-2 variant S glycoproteins. Domains independently colored corresponding to each variant including: NTD, N-terminal domain; RBD, receptor binding domain; FP, fusion peptide; HR1, heptad repeat 1; CH, central helix region; CD, connector domain; HR2, heptad repeat 2; GSAS, mutated furin cleavage site. Glycans depicted as black forks and stabilizing proline mutations colored orange. Mutations (compared with engineered Wuhan-hu-1 strain) color matched to the domains of each variant, with conserved mutations between variants labelled in red. Deleterious mutations highlighted using dashed lines.

Here, we demonstrate a broad similarity in the processing state of the glycan shield of SARS-CoV-2 VOC S proteins relative to the parental Wuhan lineage. Despite discrete differences in glycan processing across conserved N-linked glycan sites, the novel P.1 PNGSs at N20 and N188, are of interest due to their influence on local glycan processing. To better understand how the changing dynamics of the glycan shield may impact the antigenic surface of S protein, we performed molecular dynamics simulations of P.1 S glycoprotein using the site-specific glycosylation data generated by liquid chromatography-mass spectrometry. These data suggest a possible evasion mechanism through the introduction of N188, which is added into a region of the protein which has been shown to mediate the binding of a ligand, biliverdin, a metabolite involved in heme catabolism [62]. Furthermore, analysis of the B.1.351 and B.1.617.2 strains demonstrate little variability in the glycan shield. This suggests that whilst the glycan shield of SARS-CoV-2 S protein so far has proven resistant to changes, some VOCs have already demonstrated the potential for utilizing changes in the glycan shield for immune evasion, and therefore must be closely monitored in the future.

## Results and discussion

### Expression and purification of SARS-CoV-2 variant S glycoproteins

To interrogate the glycan shield of SARS-CoV-2 VOC S protein and investigate the impact on ACE2 binding, the sequences for the B.1.351 (Beta), P.1 (Gamma) and B.1.617.2 (Delta) variants were engineered to produce soluble recombinant native-like trimers. As first demonstrated by C. Hsieh, et al. 2020 [61], six stabilizing proline mutations at positions F817P, A892P, A899P, A942P, K986P, V987P were introduced, and the C-terminus of the protein was truncated at position 1208. Additionally, a C-terminal foldon domain was inserted to promote trimerization, followed by an octa-his tag to aid purification (**Figure 2**). All SARS-CoV-2 S glycoproteins were expressed in HEK 293F cells and purified using immobilized metal affinity chromatography (IMAC) followed by size exclusion chromatography (SEC) (**Supplemental Figure 1A**). For each S protein variant, a single peak was observed, which corresponds to trimeric material. This peak was collected, yielding >2 mg trimer per 200 mL expression. When subjected to SDS-PAGE, a single band between 160-200 kDa was observed, corresponding to a single protomer.

### Determining the glycan shield of SARS-CoV-2 variants using LC-MS

Using liquid chromatography-mass spectrometry (LC-MS), we explored the compositional differences in the glycan shield of SARS-CoV-2 VOCs at the site-specific level. To generate glycopeptides containing a single PNGS, trypsin, chymotrypsin, and alpha-lytic protease were utilized, and the glycopeptide products were subjected to higher energy collision induced dissociation (HCD) fragmentation. Overall, this analysis revealed a broad consensus in glycosylation at the majority of sites between the variants and the parental Wuhan lineage (**Figure 3** and **Supplemental Table 1**) [63]. All the S glycoprotein variants produced in this study displayed oligomannose-type glycan signatures at the sterically protected site, N234, suggesting that an archetypical trimeric association has been achieved (**Figure 3** and **Supplemental Tables 1-4)** [64]. As discussed by Allen et al., 2021, and Brun et al., 2021, viral derived- and recombinantly expressed S glycoprotein displayed immature glycan processing N234, suggesting a conserved quaternary protein structure influence on glycan processing at this position [52,65]. The glycosylation profile at N122 is also extremely conserved between the VOCs and the Wuhan-hu-1 lineage, presenting a heterogenous mix of complex-, hybrid- and oligomannose-type N-linked glycans. Such a mixture of different glycan processing states may be attributed to the site’s limited susceptibility to processing in the instance of poor substrate availability. To a lesser extent, similar microheterogeneity is also observed at N616 in B.1.351 (Beta), P.1 (Gamma) and B.1.617.2 (Delta). Additionally, an abundance of complex-type glycans are observed at RBD sites N331 and N343 across all SARS-CoV-2 variants, suggesting that additional structural changes arising from RBD mutations K417N/T, E484K and N501Y do not substantially impact RBD glycan processing. As observed at N74 in the prefusion stabilized recombinant Wuhan-hu-1 S protein [63], a small proportion (4-7%) of glycoforms at this position within the variants contain sulfate moieties (**Supplemental Table 1**). Interestingly, N165 glycosylation on membrane-bound S protein, both viral-derived and cell surface expressed, predominantly present complex-type glycans with high levels of galactosylation [66]. Perhaps an artifact from introducing stabilizing mutations into recombinant material, N165 presents more oligomannose-type glycoforms throughout a range of lab-expressed material, including those in the ‘HexaPro’ format [63]. Despite this, it is interesting to observe the same glycan signatures at these sites within the stabilized VOC S proteins, too. Across the trimer of every analyzed sample, complex type glycans contained high levels of fucosylation largely owing to the producer cell line, HEK 293F, while low levels of sialic acid (<50%) was observed across most complex sites, except at C-terminal sites (>70%).

**Figure 3.**
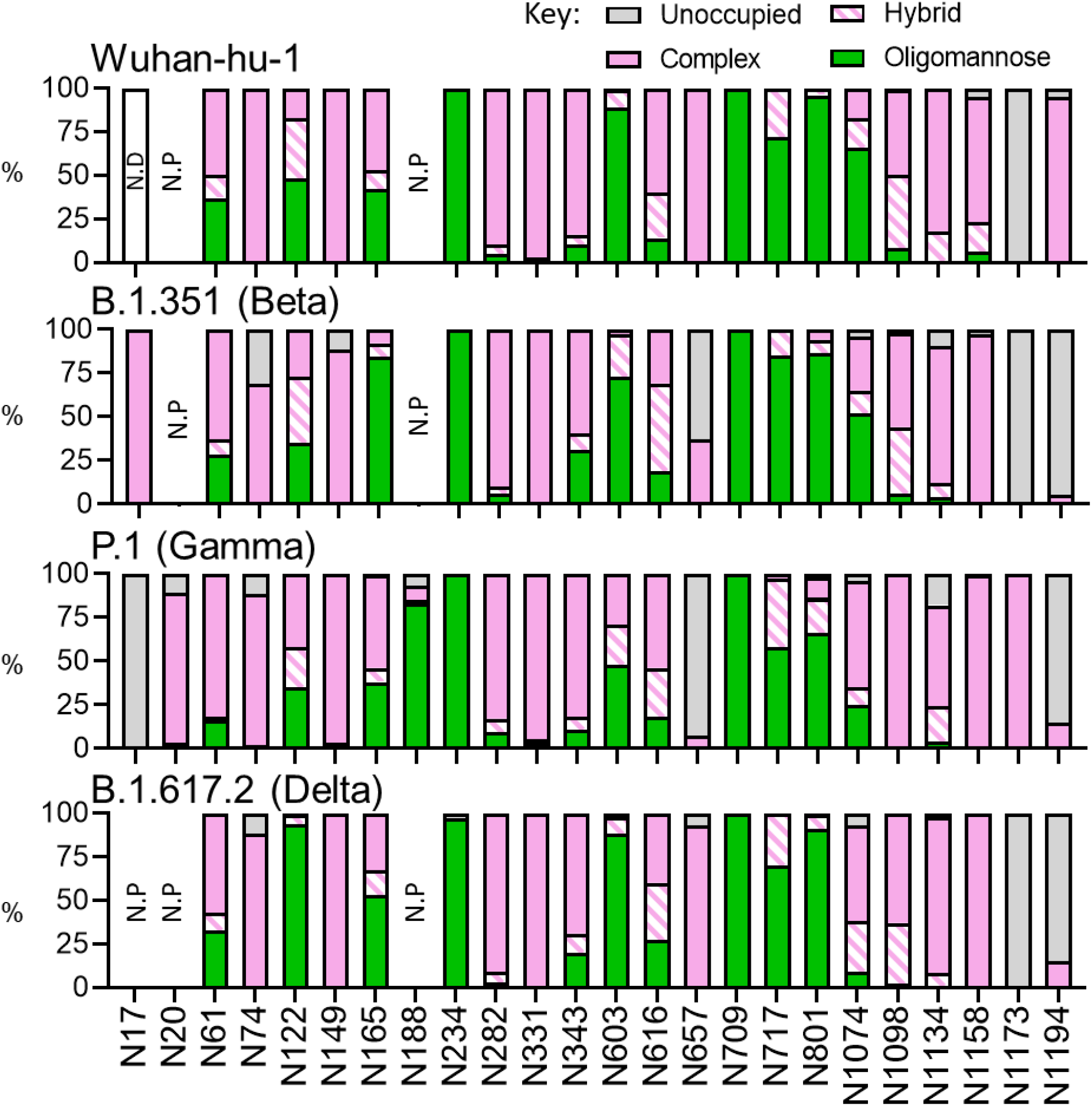
Site-specific glycan analysis of recombinant Wuhan-hu-1, B.1.351 and P.1 S protein using LC-MS. **A)** Glycan compositions are grouped into their corresponding categories, with complex-type glycans displayed in pink, hybrid as hatched pink and white, oligomannose in green and unoccupied in grey. Glycan sites not present are denoted as “N.P.” and glycan sites that could not be resolved are denoted as “N.D.”.

Whilst many sites are processed analogously to the Wuhan-hu-1 strain, there are several site-specific differences in glycosylation between the variants. The two additional N-glycan sites within the P.1 S protein exhibit disproportionate effects on adjacent glycan presentation. One of these novel sites, N20, is introduced proximal to the N17 site found in every investigated lineage (except B.1.617.2) and is likely the factor diminishing N17 glycan occupancy (**Figure 3**). Glycosylation efficiency is greatly impacted at regions of high PGNS density, like that between N17 and N20 (<u>N</u>FT<u>N</u>RT), whereby sequon skipping is expected [67]. Differing only by the second position residue, both sequons follow an N-X-T consensus yet N20 glycosylation efficiency is profoundly favored over that at N17. This may be explained by the negative bias of OST isoforms towards large hydrophobic and negatively charged side chains of the X residue in the N-X-T sequon [68]. Interestingly, the additional PNGS in P.1, N188, seems to be situated within an area of high steric protection, like N234, as reflected by the high abundance of oligomannose-type glycans detected at this site. Restricted accessibility to immature glycans by ER and Golgi mannosidases can be influenced by the local 3D protein architecture and glycan clustering, giving rise to a large abundance of immature Hex_9_GlcNAc_2_ glycoforms at enzyme-resistant sites [51,69]. Unlike N234, however, the degree of mannose trimming at N188 is more extensive (**Figure 4** and **Supplemental Table 3**), giving rise to abundance of Man_5_GlcNAc_2_, suggesting a subtle enhancement in susceptibility to *cis*-Golgi mannosidases. The P.1 lineage is the only VOC to date that has introduced a novel glycan site. The accruement of PNGS in HIV-1 Env and Influenza HA during chronic infection and endemic circulation, respectively, is retable. For this reason, the possible functional mechanisms of N188 in SARS-CoV-2 P.1 S protein warrant investigation.

**Figure 4:**
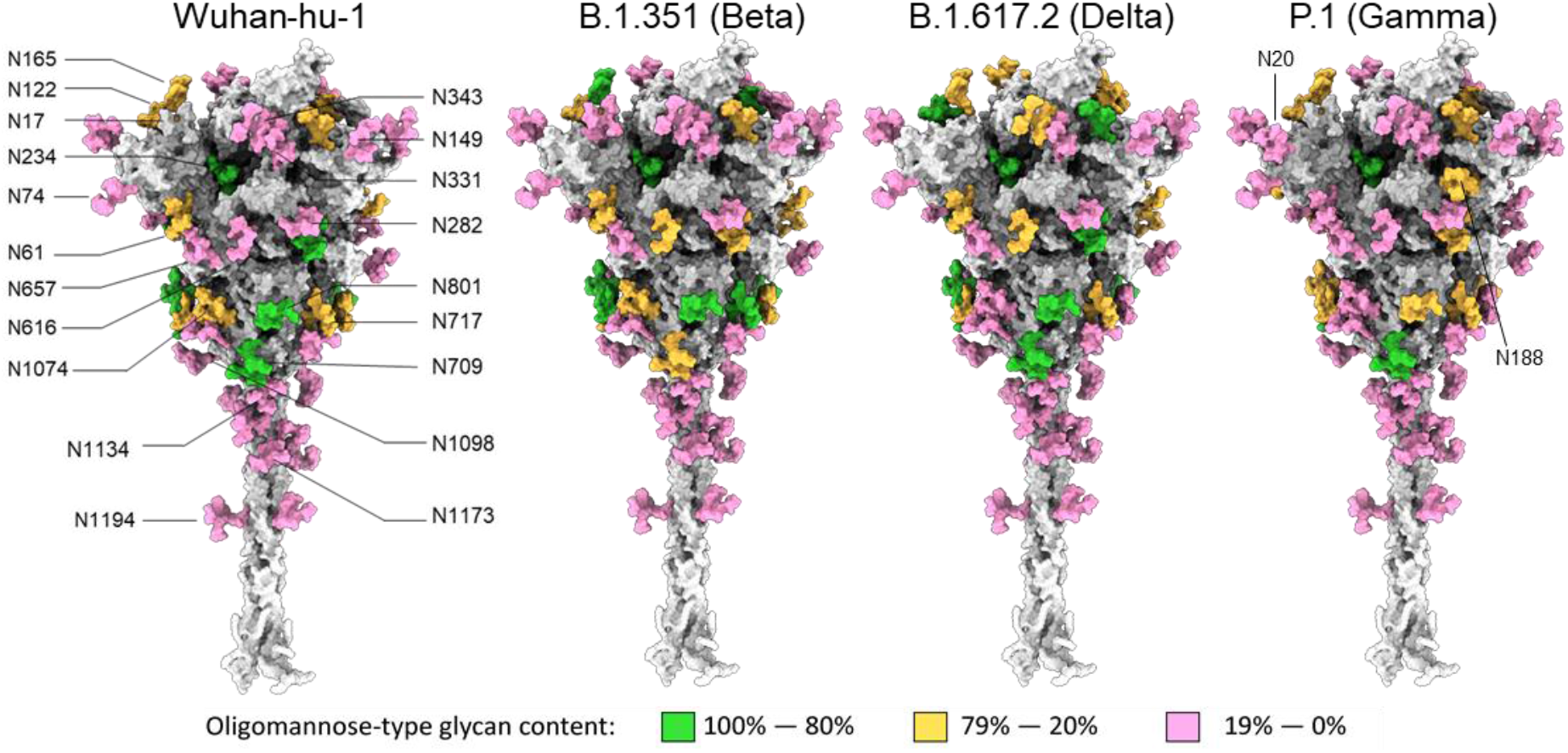
Map of the site-specific glycosylation of the SARS-CoV-2 S protein. Using a previously generated model of the Wuhan hu1 glycan shield by Zuzic et al. 2022 [34], the site-specific glycosylation determined in Figure 3 was used to color the map. If the oligomannose-type glycan content was 80% or more then the site is colored green. Between 79% and 20% correspond to mixed sites and are colored orange, and sites containing less than 20% oligomannose-type glycans correspond to complex-type glycan sites and are colored pink. Each variant is based on the Wuhan model; however, the additional glycan sites present in the Gamma variant have been added and are labelled appropriately.

### Interrogating RBD dynamics of N188-containing P.1 (Gamma) S protein

To understand how the protein landscape may shape N188 glycosylation of the P.1 (Gamma) S glycoprotein, we performed extensive sampling through conventional molecular dynamics (MD) of two P.1 S models, one bearing Man_5_GlcNAc_2_ (Man5) at N188, and another lacking glycosylation at this site. We reconstructed the P.1 S glycoprotein ectodomain from the cryo-EM structure (PDB 7SBS) [70] with a glycosylation profile matching the site-specific glycan data shown in **Figure 5**, and with a Man5 at N188 in all three protomers (**Supplemental Table 3**). The results show that for the entirety of the 1.05 μs molecular dynamics (MD) production run the N188-Man5 occupies a deep cavity in all three protomers (**Figure 5A** and **B)**, with the core and the 1-3 arm residues engaging in dispersion and hydrogen bonding interactions with different residues lining the interior of the cavity, and the loop above it (residues 171 to 183). To note, the imidazole sidechain of His207 stacks with the N188-linked GlcNAc in all three protomers and continues throughout the production trajectory. N188-Man5 can also engage with different groups of residues in the pocket, e.g., most frequently with Ser190, Asp178, Phe175 and Tyr 170, depending on its conformation (**Figure 5C**).

**Figure 5.**
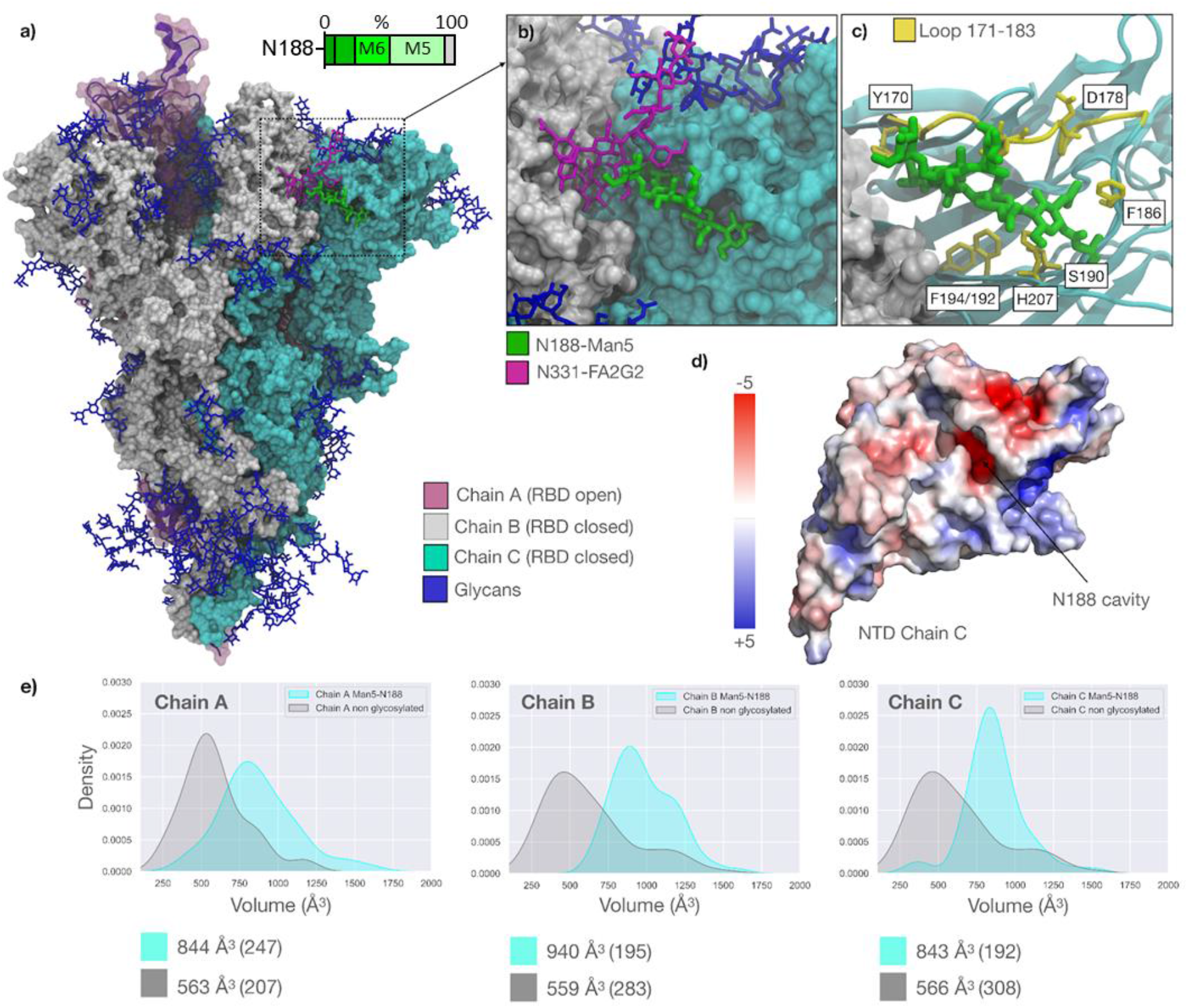
**A)** Representative snapshot (793 ns) from 1.050 μs MD production trajectory of the reconstructed SARS-CoV-2 P.1 S ectodomain from PDB 7sbs. The three protomers are color-coded according to the legend. All glycan structures are shown in blue with sticks, except for N331-FA2G2 (Chain B) and N188-Man5 (Chain C) highlighted in purple and green, respectively. Bar chart depicts the degree of oligomannose-type glycan trimming (Man_9_GlcNAc_2_-Man_5_GlcNAc_2_ denoted by M9-M5) at sites N188 and N234 in P.1 with oligomannose glycoforms displayed using increasing luminance of green. **B)** Close-up view of the NTD of Chain C with the N188-Man5 embedded in the cavity and interacting with N331-FA2G2 from the adjacent, closed RBD (Chain B). **C)** Close-up view of the NTD of Chain C rendered with cartoons (cyan), with the adjacent RBD (Chain B) as a white solvent-accessible surface. The residues that the N188-Man5 makes stable contact through the trajectory (as seen for all protomers) are highlighted with sticks (yellow) and labelled based on the PDB 7sbs numbering. The loop from residue 171 to 183 framing one side of the cavity is also highlighted in yellow. **D)** Structure of the NTD (Chain C) represented by its electrostatic potential energy surface calculated with the Adaptive Poisson-Boltzmann Solver (APBS) plugin in Pymol (www.pymol.org). The location of the cavity occupied by the N188-Man5 is indicated with an arrow. **E)** Kernel-density estimate (KDE) plots of the molecular volumes (in Å3) of the NTD cavity in all different protomers calculated from 300 ns to 1.05 μs of the MD production trajectories for the N188-glycosylated model gamma S (cyan) and for the N188-non-glycosylated model (grey). The data from 0 to 300 ns suggested a significant readjustment of the structure and was considered conformational equilibration. Median values for all distributions are indicated below the plots with corresponding standard deviations in brackets. Molecular volumes calculated with PyVol [73], plots with seaborn (https://seaborn.pydata.org/), all molecular rendering, except for the electrostatic potential surface in panel d) with VMD (https://www.ks.uiuc.edu/Research/vmd/).

The cavity occupied by the N188-Man5 in the SARS-CoV-2 S NTD is located between two β-sheets and is rich in aliphatic and aromatic residues that promote the adhesion between the sheets, and thus the stability of the domain. Asp121 and Glu96 are the only acidic residues located at the back of the pocket, (**Figure 5D**). The cavity’s electrostatic environment is well complemented by the amphipathic nature of the N-glycan, more so than by water. Indeed, in the absence of the N188 glycan, assessed through an additional 1.05 μs MD trajectory of the N188-non glycosylated P.1 S model, the volume of the cavity is significantly reduced (**Figure 5E**). This suggests a partial collapse of the cavity or a reduced accessibility. This effect is particularly pronounced in the P.1 S NTDs because of the R190S mutation, which introduces the sequon at N188. Previous MD simulations of the Wuhan-hu-1 S protein [71,72] show that the Arg190 can form a salt-bridge with Asp178 (**Figure 5C**), which widens and possibly facilitates access to the cavity. Indeed, the same NTD cavity has been shown to bind the heme metabolites bilirubin and biliverdin [62], with important consequences to antibody immunity. Structural studies of the SARS-CoV-2 S glycoprotein have shown that biliverdin enters the SARS-CoV-2 (Wuhan-hu-1) S NTD cavity deeper than the N188-Man5 does in the P.1 S (**Supplemental Figure 2A**) [62], yet it forms contacts with some of the same residues, such as His207, which can form stacking interactions with both molecules. A complete set of binding assays show that the binding of biliverdin significantly decreases recognition of the Wuhan-hu-1 S by 17% of the 53 IgGs in the panel, strongly indicating a reduced immune recognition against the S-biliverdin complex [62]. This effect was linked to a conformational change of the loop comprising residues 173 to 188 occurring upon binding of biliverdin, which is a known epitope recognized by a subset of NTD-specific antibodies [74]. The binding of biliverdin shifts the loop from an ‘open’ to a ‘closed’ conformation, the latter of which is not recognized, (**Supplemental Figure 2B**). Within this framework, the MD simulations of the P.1 S we performed show that the presence of the N188-Man5 in the cavity displays a similar shift, whereby the loop samples conformations analogous to the ‘closed’ conformation observed in the presence of biliverdin (PDB 7NTA) (**Supplemental Figure 2B**). From these simulations, it is reasonable to infer that the appearance of a new sequon at N188 may have contributed to the higher infectivity of the SARS-CoV-2 P.1 relative to Wuhan hu-1, with the N188 glycan shielding the protein surface in an analogous way to the binding of heme metabolites. These studies demonstrate the potential for additional immune evasion mechanisms beyond simple changes in the amino acid sequence of the protein and highlight the importance of understanding the structure and function of the SARS-CoV-2 glycan shield.

### Perspectives

In this study, we sought to understand potential immune evasion mechanisms involving changes in the glycan shield. By combining compositional glycan analysis with molecular dynamics simulations, we provide molecular insights into how the P.1 (Gamma) variant with the additional N188 glycan site exhibits increased shielding of the surface of the spike glycoprotein. In addition, we show limited changes in the glycan shield of other key VOCs, perhaps suggesting that SARS-CoV-2 is yet to fully exploit the potential of glycan-mediated immune evasion, as other viruses have. Despite containing over sixty N-glycan sites across the trimer, the glycan shield density of SARS-CoV-2 S protein is sparse relative to that of HIV-1, Influenza and LASV, leaving much of the immunogenic protein surface exposed to the humoral immune system [75]. As a virus that has only recently begun circulating in the human population, SARS-CoV-2 has so far exploited more simple adaptations to improve infectivity and evade host immunity, such as changes in the RBD. Modulating the glycan shield likely has detrimental impacts on viral infectivity. As observed in Influenza H3N2, periodic endemic circulation and evolving immunity within the human population has contributed to the accumulation of N-linked glycan sites on the haemagglutinin ectodomain, and a consequential increase in oligomannose-type glycan clustering. The eventual plateau of PNGS accruement suggests a fitness cost associated with increased glycan shielding[58]. In fact, relative to other coronaviruses, SARS-CoV-2 lacks a glycosylation site in the RBD, N370, and this has been suggested to enhance infectivity of SARS-CoV-2 [63]. Perhaps, as vaccinations and infections continue, viral evolution will exhaust simple amino acid substitutions and deletions to escape the evolving immunity of the human population and begin to adapt the glycan shield, with the potential fitness costs this may involve. Other coronaviruses that have been circulating in the human population for longer, such as the common cold coronavirus, NL63, are much more densely glycosylated, containing nearly double (40) the number of glycan sites per protomer as for SARS-CoV-2 [48]. Therefore, tools such as those described in this manuscript, will assist in exploring how the glycans of viral spike glycoproteins are being used to subvert the host immune system, as new variants of concern emerge.

## Materials and methods

### Design of prefusion-stabilized soluble SARS-CoV-2 S variants

All engineered SARS-CoV-2 S variant ectodomains were designed based on the HexaPro gene (Addgene plasmid #154754) containing stabilizing proline mutations at positions F817P, A892P, A899P, A942P, K986P, V987P and mutated furin cleavage site (GSAS) at residues 682-685. Constructs were truncated at residue 1213 upstream of the transmembrane region and a T4 foldon domain (GYIPEAPRDGQAYVRKDGEWVLLSTFL) introduced at the C-terminus of the protein to promote trimerization. To assist purification, a HRV3C protease site followed by an octa-His tag was added to the C-terminus of the gene and cloned into the pαH expression vector.

### Expression and purification of SARS-CoV-2 variants

HEK 293F cells were transiently transfected with the mammalian expression plasmid pαH containing SARS-CoV-2 variant S ectodomains. 293F cells were cultured in FreeStyle 293 Expression Medium (Fisher Scientific) and maintained at a density of 0.2-3 × 10^6^ cells/mL at 37°C, 8% CO_2_ and 125 rpm shaking. Prior to transfection, two solutions of 25 mL Opti-MEM (Fisher Scientific) medium were prepared. Expression plasmid encoding SARS-CoV-2 S glycoprotein was added to the first solution to give a final concentration of 310 μg/L. To the other solution, 1 mg/mL pH7 polyethylenimine (PEI) max reagent was added to generate a ratio of 3:1 PEI max:plasmid DNA. Both solutions were combined and incubated for 30 minutes at room temperature. Cells were transfected at a density of 1 × 10^6^ cells/mL and incubated for 7 days at 37°C, 8% CO_2_ and 125 rpm shaking.

Cells were centrifuged at 3041 × g for 30 minutes at 4°C and supernatant was applied to a 500 mL Stericup-HV sterile vacuum filtration system (Merck) with a pore size of 0.22 µm. Purification of SARS-CoV-2 S glycoprotein was undertaken using an ÄKTA Pure system (Cytiva). A 5 mL HisTrap Excel column (Cytiva) charged with Ni(II) was equilibrated using 10 column volumes (CV) of washing buffer (50 mM Na_2_PO_4_, 300 mM NaCl) at pH 7. Supernatant was then loaded onto the column at a flow rate of 5 mL/min and washed with 10 column volumes (CV) of washing buffer containing 50 mM imidazole. Protein was eluted from the column in 3 CV of elution buffer (300 mM imidazole in washing buffer) and buffer exchanged to phosphate buffered saline (PBS) and concentrated using a Vivaspin column (MWCO 100 kDa) (Cytiva).

The nickel purified eluate was concentrated to 1 mL in PBS and injected into a Superdex 200 pg 16/600 column (Cytiva) to further purify trimeric S glycoprotein using size exclusion chromatography (SEC). The column was washed with PBS at 1 mL/min for 2 hours where fractions corresponding to the correct peak on the size exclusion chromatogram were collected and concentrated to ∼1 mL as above.

### Mass spectrometry of glycopeptides

SARS-CoV-2 variant S glycoproteins were denatured for 1h in 50 mM Tris/HCl, pH 8.0 containing 6 M of urea. Next, the sample was reduced and alkylated by adding 5 mM dithiothreitol (DTT) and 20 mM iodoacetamide (IAA) and incubated for 1h in the dark, followed by a 1h incubation with 20 mM DTT to eliminate residual IAA. The alkylated S glycoproteins were buffer exchanged into 50 mM Tris/HCl, pH 8.0 using Vivaspin columns (3 kDa) and two of the aliquots were digested separately overnight using trypsin (Mass Spectrometry grade, Promega), chymotrypsin (Mass Spectrometry Grade, Promega) or alpha lytic protease (Sigma Aldrich) at a ratio of 1:30 (w/w) at 37°C. The next day, the peptides were dried and extracted using C18 Zip-tip (MerckMilipore). The peptides were dried again, re-suspended in 0.1% formic acid and analyzed by nanoLC-ESI MS with an Ultimate 3000 HPLC (Thermo Fisher Scientific) system coupled to an Orbitrap Eclipse mass spectrometer (Thermo Fisher Scientific) using stepped higher energy collision-induced dissociation (HCD) fragmentation. Peptides were separated using an EasySpray PepMap RSLC C18 column (75 µm × 75 cm). A trapping column (PepMap 100 C18 3μm particle size, 75μm × 2cm) was used in line with the LC prior to separation with the analytical column. The LC conditions were as follows: 280-minute linear gradient consisting of 4-32% acetonitrile in 0.1% formic acid over 260 minutes followed by 20 minutes of alternating 76% acetonitrile in 0.1% formic acid and 4% ACN in 0.1% formic acid, used to ensure all the sample had eluted from the column. The flow rate was set to 300 nL/min. The spray voltage was set to 2.5 kV and the temperature of the heated capillary was set to 40 °C. The ion transfer tube temperature was set to 275 °C. The scan range was 375−1500 m/z. Stepped HCD collision energy was set to 15, 25 and 45% and the MS2 for each energy was combined. Precursor and fragment detection were performed using an Orbitrap at a resolution MS1= 120,000. MS2= 30,000. The AGC target for MS1 was set to standard and injection time set to auto which involves the system setting the two parameters to maximize sensitivity while maintaining cycle time. Full LC and MS methodology can be extracted from the appropriate Raw file using XCalibur FreeStyle software or upon request.

Glycopeptide fragmentation data were extracted from the raw file using Byos (Version 4.0; Protein Metrics Inc.). The glycopeptide fragmentation data were evaluated manually for each glycopeptide; the peptide was scored as true-positive when the correct b and y fragment ions were observed along with oxonium ions corresponding to the glycan identified. The MS data was searched using the Protein Metrics 305 N-glycan library with sulfated glycans added manually. The relative amounts of each glycan at each site as well as the unoccupied proportion were determined by comparing the extracted chromatographic areas for different glycotypes with an identical peptide sequence. All charge states for a single glycopeptide were summed. The precursor mass tolerance was set at 4 ppm and 10 ppm for fragments. A 1% false discovery rate (FDR) was applied. The relative amounts of each glycan at each site as well as the unoccupied proportion were determined by comparing the extracted ion chromatographic areas for different glycopeptides with an identical peptide sequence. Glycans were categorized according to the composition detected. HexNAc(2), Hex(9−3) was classified as M9 to M3. Any of these compositions that were detected with a fucose are classified as FM. HexNAc(3)Hex(5−6)Neu5Ac)(0-4) was classified as Hybrid with HexNAc(3)Hex(5-6)Fuc(1)NeuAc(0-1) classified as Fhybrid. Complex-type glycans were classified according to the number of processed antenna and fucosylation. Complex glycans are categorized as HexNAc(3)(X), HexNAc(3)(F)(X), HexNAc(4)(X), HexNAc(4)(F)(X), HexNAc(5)(X), HexNAc(5)(F)(X), HexNAc(6+)(X) and HexNAc(6+)(F)(X). Core glycans are any glycan smaller than HexNAc(2)Hex(3).

### Computational Methods

The model of the P.1 S glycoprotein ectodomain was built from the cryo-EM structure (PDB 7SBS) [70] with a glycosylation profile matching the site-specific glycan data in this work. The effect of the N188 glycosylation was determined by analyzing the dynamics of two systems, one with a Man5 at N188 in all three protomers and one without the N-glycan at N188 in all three protomers. Each simulation performed three times through uncorrelated production runs for each model system. In all MD simulations the protein and counterions (200 mM) were represented by the AMBER ff14SB parameter set [76], whereas the glycans were represented by the GLYCAM06j-1 version of the GLYCAM06 force field [77]. Water molecules were represented by the TIP3P model. All simulations were run with v18 of the AMBER software package [78]. The following running protocol was used for all MD simulations. The energy of the S ectodomain models was minimized in two steps of 50 k cycles of the steepest descent algorithm each. During the first minimization all the heavy atoms were kept harmonically restrained using a potential weight of 5 kcal mol^−1^Å^−2^, while the solvent, counterions and hydrogen atoms were left unrestrained. The minimization step was repeated with only the protein heavy atoms restrained. After energy minimization the system was equilibrated in the NVT ensemble with the same restraints scheme, where heating was performed in two stages over a total time of 1 ns, from 0 to 100 K (stage 1) and then from 100 to 300 K (stage 2). During equilibration the SHAKE algorithm was used to constrain all bonds to hydrogen atoms. The Van der Waals interactions were truncated at 11 Å and Particle Mesh Ewald (PME) was used to treat long range electrostatics with B-spline interpolation of order 4. Langevin dynamics with collision frequency of 1.0 ps^−1^ was used to control temperature, with a pseudo-random variable seed to ensure there are no synchronization artefacts. Once the system was brought to 300 K an equilibration phase in the NPT ensemble of 1 ns was used to set the pressure to 1 atm. The pressure was held constant with isotropic pressure scaling and a pressure relaxation time of 2.0 ps. At this point all restraints on the protein heavy atoms were removed, allowing the system to evolve for 15 ns of conformational equilibration before production of 1.05 µs for each replica, generated from different starting velocities. The MD simulations were performed on resources from the Irish Centre for High-End Computing (www.ichec.ie). Structures in PDB format are deposited on https://github.com/CFogarty-2275/Gamma_Spike.

## Authorship contribution

**Maddy L. Newby:** Conceptualization, Investigation, Formal analysis, Writing – Original Draft. **Joel D. Allen:** Investigation, Formal Analysis, Writing – Original Draft. **Carl A. Fogarty:** Investigation, Formal Analysis, Writing – Original Draft. **John Butler:** Investigation. **Elisa Fadda:** Investigation, Funding Acquisition, Formal Analysis, Writing – Original Draft. **Max Crispin:** Conceptualization, Funding Acquisition, Supervision, Writing – Original Draft.

## Acknowledgements

This work was supported by the International AIDS Vaccine Initiative (IAVI) through grant INV-008352/OPP1153692 funded by the Bill and Melinda Gates Foundation (M.C.). We also gratefully acknowledge support from the University of Southampton Coronavirus Response Fund (M.C.), and a donation from the Bright Future Trust (M.C.). The Science Foundation of Ireland (20/FFP-P/8809) is gratefully acknowledged for financial support (E.F.). The Irish Research Council is gratefully acknowledged for funding under the Government of Ireland Postgraduate Scholarship scheme (C.F.). The Irish Centre for High-End Computing (www.ichec.ie) is gratefully acknowledged for the generous allocation of computational resources (C.F. and E.F.). The opinions, findings and conclusions or recommendations expressed in this material are those of the author(s) and do not necessarily reflect the views of the Science Foundation Ireland.

## Notes

### Competing Interest Statement

The authors have declared no competing interest.

