## Supplemental information for "Natural variations within the glycan shield of SARS-CoV-2 impact viral spike dynamics"

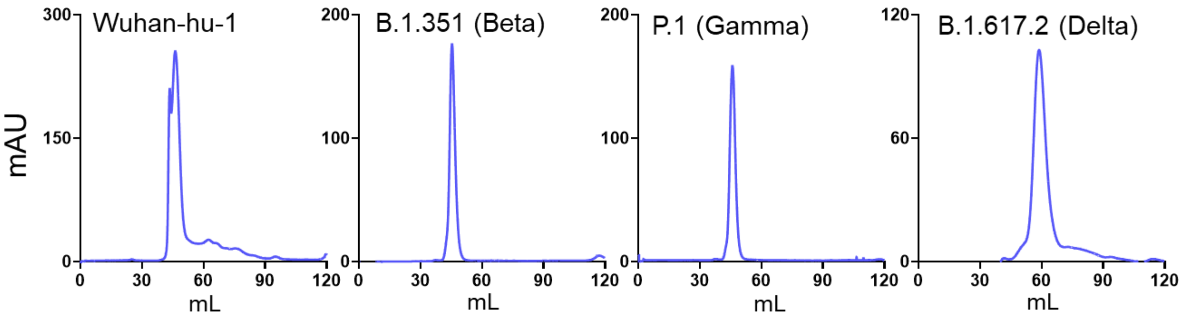


**Supplemental Figure 1– Size Exclusion Chromatographs of SARS-CoV-2 variants.** (A) Size exclusion chromatogram of Wuhan-hu-1, B.1.351, P.1, and B.1.617.2 lineages following nickel affinity chromatography. SDS-PAGE using Coomassie blue stain of pooled SEC fractions corresponding to trimeric SARS-CoV-2 S glycoprotein.

### Supplemental Table 1: Site-specific glycan analysis of Wuhan hu1 spike glycoprotein

| | N17 | N61 | N74 | N122 | N149 | N165 | N234 | N282 | N331 | N343 | N603 | N616 | N657 | N709 | N717 | N801 | N1074 | N1098 | N1134 | N1158 | N1173 | N1194 |
|---|---|---|---|---|---|---|---|---|---|---|---|---|---|---|---|---|---|---|---|---|---|---|
| M9Glc | | 0 | 0 | 0 | 0 | 0 | 0 | 0 | 0 | 0 | 0 | 0 | 0 | 0 | 0 | 0 | 0 | 0 | 0 | 0 | 0 | 0 |
| M9 | | 0 | 0 | 0 | 0 | 0 | 72 | 0 | 0 | 0 | 0 | 0 | 0 | 0 | 0 | 0 | 0 | 0 | 0 | 0 | 0 | 0 |
| M8 | | 0 | 0 | 0 | 0 | 0 | 16 | 1 | 0 | 0 | 2 | 0 | 0 | 0 | 0 | 6 | 1 | 0 | 0 | 1 | 0 | 0 |
| M7 | | 0 | 0 | 5 | 0 | 4 | 5 | 0 | 0 | 0 | 9 | 0 | 0 | 37 | 3 | 4 | 1 | 2 | 0 | 0 | 0 | 0 |
| M6 | | 1 | 0 | 3 | 0 | 4 | 3 | 0 | 0 | 0 | 9 | 0 | 0 | 63 | 13 | 16 | 4 | 1 | 0 | 0 | 0 | 0 |
| M5 | | 30 | 0 | 39 | 1 | 33 | 4 | 4 | 2 | 8 | 44 | 13 | 0 | 0 | 48 | 63 | 58 | 3 | 1 | 4 | 0 | 2 |
| M4 | | 4 | 0 | 1 | 0 | 1 | 0 | 0 | 0 | 1 | 22 | 0 | 0 | 0 | 8 | 6 | 0 | 0 | 0 | 0 | 0 | 0 |
| M3 | | 1 | 0 | 0 | 0 | 0 | 0 | 0 | 0 | 0 | 2 | 0 | 0 | 0 | 0 | 1 | 0 | 0 | 0 | 0 | 0 | 0 |
| FM | | 0 | 0 | 0 | 0 | 0 | 0 | 0 | 0 | 0 | 0 | 0 | 0 | 0 | 0 | 0 | 0 | 0 | 0 | 0 | 0 | 0 |
| Hybrid | | 10 | 0 | 30 | 0 | 9 | 0 | 4 | 0 | 2 | 10 | 17 | 0 | 0 | 28 | 4 | 12 | 34 | 1 | 6 | 0 | 0 |
| Fhybrid | | 2 | 0 | 5 | 0 | 2 | 0 | 1 | 0 | 5 | 0 | 9 | 0 | 0 | 0 | 0 | 8 | 9 | 16 | 11 | 0 | 0 |
| HexNAc(3)(x) | | 4 | 0 | 4 | 0 | 2 | 0 | 4 | 0 | 1 | 1 | 8 | 0 | 0 | 0 | 0 | 0 | 4 | 0 | 2 | 0 | 0 |
| HexNAc(3)(F)(x) | | 2 | 1 | 2 | 2 | 1 | 0 | 0 | 1 | 5 | 0 | 3 | 0 | 0 | 0 | 0 | 3 | 3 | 7 | 8 | 0 | 0 |
| HexNAc(4)(x) | | 24 | 0 | 6 | 0 | 15 | 0 | 20 | 0 | 0 | 0 | 23 | 0 | 0 | 0 | 0 | 2 | 17 | 0 | 5 | 0 | 0 |
| HexNAc(4)(F)(x) | | 7 | 8 | 4 | 50 | 8 | 0 | 8 | 53 | 49 | 0 | 12 | 13 | 0 | 0 | 0 | 8 | 15 | 53 | 36 | 0 | 4 |
| HexNAc(5)(x) | | 11 | 0 | 0 | 0 | 12 | 0 | 34 | 0 | 0 | 0 | 14 | 0 | 0 | 0 | 0 | 0 | 2 | 0 | 0 | 0 | 0 |
| HexNAc(5)(F)(x) | | 2 | 83 | 1 | 45 | 8 | 0 | 17 | 42 | 28 | 0 | 0 | 83 | 0 | 0 | 0 | 4 | 5 | 19 | 15 | 0 | 10 |
| HexNAc(6+)(x) | | 1 | 0 | 0 | 0 | 0 | 0 | 5 | 0 | 0 | 0 | 0 | 0 | 0 | 0 | 0 | 0 | 1 | 0 | 0 | 0 | 0 |
| HexNAc(6+)(F)(x) | | 0 | 8 | 0 | 1 | 0 | 0 | 2 | 0 | 0 | 0 | 0 | 4 | 0 | 0 | 0 | 0 | 1 | 3 | 5 | 0 | 79 |
| Unoccupied | | 0 | 0 | 0 | 0 | 0 | 0 | 0 | 0 | 0 | 0 | 0 | 0 | 0 | 0 | 0 | 0 | 1 | 0 | 5 | 100 | 5 |
| Core | | 0 | 0 | 0 | 0 | 0 | 0 | 0 | 1 | 0 | 1 | 0 | 0 | 0 | 0 | 0 | 0 | 0 | 0 | 0 | 0 | 0 |
| Oligomannose | | 37 | 0 | 48 | 1 | 42 | 100 | 5 | 2 | 10 | 88 | 14 | 0 | 100 | 72 | 96 | 63 | 8 | 1 | 6 | 0 | 2 |
| Hybrid | | 13 | 0 | 35 | 0 | 11 | 0 | 4 | 0 | 6 | 10 | 26 | 0 | 0 | 28 | 4 | 19 | 42 | 17 | 17 | 0 | 0 |
| Complex | | 50 | 100 | 17 | 99 | 47 | 0 | 90 | 97 | 83 | 1 | 60 | 100 | 0 | 0 | 0 | 17 | 49 | 82 | 72 | 0 | 94 |
| Unoccupied | | 0 | 0 | 0 | 0 | 0 | 0 | 0 | 0 | 0 | 0 | 0 | 0 | 0 | 0 | 0 | 0 | 1 | 0 | 5 | 100 | 5 |
| Fucose | | 13 | 100 | 12 | 99 | 19 | 0 | 28 | 97 | 87 | 1 | 24 | 100 | 0 | 0 | 0 | 23 | 33 | 98 | 76 | 0 | 94 |
| NeuAc | | 20 | 100 | 19 | 67 | 17 | 0 | 20 | 28 | 18 | 0 | 7 | 38 | 0 | 21 | 0 | 4 | 70 | 24 | 27 | 0 | 81 |

### Supplemental Table 2: Site-specific glycan analysis of B.1.351 (Beta) spike glycoprotein

| | N17 | N61 | N74 | N122 | N149 | N165 | N234 | N282 | N331 | N343 | N603 | N616 | N657 | N709 | N717 | N801 | N1074 | N1098 | N1134 | N1158 | N1173 | N1194 |
|---|---|---|---|---|---|---|---|---|---|---|---|---|---|---|---|---|---|---|---|---|---|---|
| M9Glc | 0 | 0 | 0 | 0 | 0 | 0 | 2 | 0 | 0 | 0 | 0 | 0 | 0 | 0 | 0 | 0 | 0 | 0 | 0 | 0 | 0 | 0 |
| M9 | 0 | 0 | 0 | 0 | 0 | 0 | 81 | 0 | 0 | 0 | 0 | 0 | 0 | 0 | 0 | 0 | 0 | 0 | 0 | 0 | 0 | 0 |
| M8 | 0 | 0 | 0 | 0 | 0 | 0 | 13 | 0 | 0 | 0 | 1 | 0 | 0 | 0 | 0 | 3 | 0 | 0 | 0 | 0 | 0 | 0 |
| M7 | 0 | 0 | 0 | 3 | 0 | 2 | 2 | 0 | 0 | 0 | 6 | 0 | 0 | 8 | 13 | 12 | 0 | 1 | 0 | 0 | 0 | 0 |
| M6 | 0 | 0 | 0 | 2 | 0 | 21 | 0 | 0 | 0 | 0 | 5 | 1 | 0 | 37 | 31 | 9 | 2 | 1 | 0 | 0 | 0 | 0 |
| M5 | 7 | 20 | 0 | 27 | 0 | 60 | 1 | 5 | 0 | 18 | 57 | 14 | 0 | 29 | 47 | 60 | 41 | 3 | 4 | 0 | 0 | 0 |
| M4 | 0 | 4 | 0 | 1 | 0 | 1 | 0 | 0 | 0 | 2 | 1 | 4 | 0 | 0 | 0 | 1 | 1 | 0 | 0 | 0 | 0 | 0 |
| M3 | 0 | 2 | 0 | 0 | 0 | 1 | 0 | 0 | 0 | 1 | 0 | 1 | 0 | 0 | 0 | 0 | 1 | 0 | 0 | 0 | 0 | 0 |
| FM | 0 | 0 | 0 | 0 | 0 | 0 | 0 | 0 | 0 | 0 | 0 | 0 | 0 | 1 | 0 | 0 | 0 | 0 | 0 | 0 | 0 | 0 |
| Hybrid | 11 | 7 | 0 | 23 | 0 | 4 | 0 | 2 | 0 | 3 | 22 | 35 | 0 | 17 | 8 | 8 | 5 | 25 | 0 | 0 | 0 | 0 |
| Fhybrid | 1 | 1 | 0 | 11 | 0 | 3 | 0 | 0 | 0 | 3 | 1 | 6 | 0 | 0 | 0 | 0 | 5 | 8 | 6 | 0 | 0 | 0 |
| HexNAc(3)(x) | 4 | 4 | 0 | 4 | 0 | 0 | 0 | 5 | 0 | 1 | 1 | 8 | 0 | 5 | 0 | 1 | 2 | 4 | 1 | 0 | 0 | 0 |
| HexNAc(3)(F)(x) | 1 | 0 | 2 | 9 | 1 | 1 | 0 | 0 | 1 | 6 | 0 | 0 | 0 | 3 | 0 | 0 | 4 | 4 | 3 | 0 | 0 | 0 |
| HexNAc(4)(x) | 13 | 25 | 0 | 3 | 0 | 1 | 0 | 27 | 0 | 1 | 0 | 15 | 0 | 0 | 0 | 1 | 3 | 18 | 3 | 0 | 0 | 0 |
| HexNAc(4)(F)(x) | 6 | 7 | 10 | 14 | 26 | 3 | 0 | 15 | 28 | 46 | 0 | 6 | 8 | 1 | 0 | 2 | 10 | 19 | 47 | 0 | 0 | 0 |
| HexNAc(5)(x) | 6 | 20 | 0 | 0 | 1 | 0 | 0 | 31 | 0 | 0 | 0 | 6 | 30 | 0 | 0 | 0 | 1 | 3 | 0 | 50 | 0 | 0 |
| HexNAc(5)(F)(x) | 50 | 8 | 44 | 2 | 59 | 2 | 0 | 12 | 20 | 18 | 0 | 3 | 9 | 0 | 0 | 0 | 7 | 6 | 15 | 1 | 0 | 1 |
| HexNAc(6+)(x) | 0 | 1 | 0 | 0 | 0 | 0 | 0 | 1 | 0 | 0 | 0 | 0 | 18 | 0 | 0 | 0 | 0 | 1 | 0 | 0 | 0 | 1 |
| HexNAc(6+)(F)(x) | 0 | 0 | 17 | 0 | 5 | 0 | 0 | 1 | 1 | 1 | 0 | 0 | 1 | 0 | 0 | 0 | 0 | 1 | 2 | 48 | 0 | 15 |
| Unoccupied | 0 | 0 | 27 | 0 | 8 | 0 | 0 | 0 | 50 | 0 | 4 | 0 | 34 | 0 | 0 | 3 | 17 | 5 | 20 | 1 | 100 | 82 |
| Core | 0 | 1 | 0 | 0 | 1 | 1 | 0 | 0 | 0 | 1 | 0 | 0 | 1 | 0 | 0 | 0 | 0 | 0 | 0 | 0 | 0 | 0 |

| | N17 | N61 | N74 | N122 | N149 | N165 | N234 | N282 | N331 | N343 | N603 | N616 | N657 | N709 | N717 | N801 | N1074 | N1098 | N1134 | N1158 | N1173 | N1194 |
|---|---|---|---|---|---|---|---|---|---|---|---|---|---|---|---|---|---|---|---|---|---|---|
| Oligomannose | 8 | 26 | 0 | 33 | 0 | 86 | 100 | 5 | 0 | 21 | 70 | 20 | 0 | 74 | 91 | 84 | 45 | 6 | 4 | 0 | 0 | 0 |
| Hybrid | 11 | 8 | 0 | 34 | 0 | 7 | 0 | 2 | 0 | 6 | 23 | 41 | 0 | 17 | 8 | 8 | 11 | 33 | 6 | 0 | 0 | 0 |
| Complex | 81 | 66 | 73 | 33 | 92 | 8 | 0 | 93 | 49 | 73 | 2 | 38 | 66 | 9 | 0 | 4 | 27 | 57 | 70 | 99 | 0 | 18 |
| Unoccupied | 0 | 0 | 27 | 0 | 8 | 0 | 0 | 0 | 50 | 0 | 4 | 0 | 34 | 0 | 0 | 3 | 17 | 5 | 20 | 1 | 100 | 82 |
| Fucose | 58 | 17 | 73 | 36 | 90 | 8 | 0 | 29 | 49 | 74 | 2 | 16 | 18 | 4 | 0 | 3 | 27 | 39 | 72 | 49 | 0 | 17 |
| NeuAc | 26 | 13 | 36 | 16 | 28 | 1 | 0 | 15 | 26 | 18 | 0 | 5 | 31 | 22 | 3 | 2 | 6 | 55 | 27 | 42 | 0 | 11 |

**Supplemental Table 3: Site-specific glycan analysis of P.1 (Gamma) spike glycoprotein**

| | N17 | N20 | N61 | N74 | N122 | N149 | N165 | N188 | N234 | N282 | N331 | N343 | N603 | N616 | N657 | N709 | N717 | N801 | N1074 | N1098 | N1134 | N1158 | N1173 | N1194 |
|---|---|---|---|---|---|---|---|---|---|---|---|---|---|---|---|---|---|---|---|---|---|---|---|---|
| M9Glc | 0 | 0 | 0 | 0 | 0 | 0 | 0 | 0 | 1 | 0 | 0 | 0 | 0 | 0 | 0 | 0 | 0 | 0 | 0 | 0 | 0 | 0 | 0 | 0 |
| M9 | 0 | 0 | 0 | 0 | 0 | 0 | 0 | 0 | 64 | 0 | 0 | 0 | 0 | 0 | 0 | 0 | 0 | 0 | 0 | 0 | 0 | 0 | 0 | 0 |
| M8 | 0 | 0 | 0 | 0 | 0 | 0 | 0 | 5 | 27 | 0 | 0 | 0 | 0 | 0 | 0 | 15 | 0 | 4 | 0 | 0 | 0 | 0 | 0 | 0 |
| M7 | 0 | 0 | 0 | 0 | 4 | 0 | 0 | 11 | 5 | 1 | 0 | 0 | 3 | 1 | 0 | 44 | 8 | 10 | 0 | 0 | 1 | 0 | 0 | 0 |
| M6 | 0 | 0 | 0 | 0 | 3 | 0 | 8 | 21 | 2 | 0 | 0 | 0 | 3 | 2 | 0 | 13 | 18 | 3 | 1 | 0 | 1 | 0 | 0 | 0 |
| M5 | 0 | 3 | 12 | 1 | 23 | 1 | 29 | 37 | 1 | 4 | 2 | 8 | 35 | 12 | 0 | 24 | 31 | 37 | 12 | 1 | 1 | 0 | 0 | 0 |
| M4 | 0 | 0 | 1 | 0 | 1 | 0 | 0 | 1 | 0 | 0 | 0 | 1 | 1 | 0 | 0 | 0 | 2 | 4 | 3 | 0 | 0 | 0 | 0 | 0 |
| M3 | 0 | 0 | 1 | 0 | 0 | 0 | 1 | 0 | 0 | 0 | 0 | 1 | 1 | 0 | 0 | 2 | 0 | 1 | 2 | 0 | 0 | 0 | 0 | 0 |
| FM | 0 | 0 | 0 | 0 | 0 | 0 | 0 | 0 | 0 | 0 | 0 | 0 | 0 | 0 | 0 | 0 | 0 | 0 | 0 | 0 | 0 | 0 | 0 | 0 |
| Hybrid | 0 | 0 | 1 | 0 | 19 | 0 | 4 | 2 | 0 | 7 | 0 | 1 | 16 | 15 | 0 | 0 | 38 | 21 | 4 | 0 | 13 | 0 | 0 | 0 |
| Fhybrid | 0 | 0 | 0 | 0 | 6 | 0 | 2 | 0 | 0 | 1 | 1 | 4 | 6 | 6 | 0 | 0 | 0 | 0 | 3 | 1 | 5 | 0 | 0 | 0 |
| HexNAc(3)(x) | 0 | 0 | 3 | 0 | 3 | 0 | 0 | 1 | 0 | 7 | 0 | 1 | 4 | 4 | 0 | 0 | 3 | 3 | 1 | 1 | 1 | 0 | 0 | 0 |
| HexNAc(3)(F)(x) | 0 | 1 | 0 | 2 | 6 | 1 | 2 | 0 | 0 | 0 | 3 | 4 | 3 | 4 | 0 | 0 | 0 | 0 | 3 | 6 | 2 | 0 | 0 | 0 |
| HexNAc(4)(x) | 0 | 0 | 59 | 0 | 8 | 0 | 8 | 1 | 0 | 39 | 0 | 0 | 2 | 12 | 0 | 0 | 0 | 8 | 1 | 0 | 15 | 0 | 0 | 0 |
| HexNAc(4)(F)(x) | 0 | 32 | 10 | 17 | 21 | 41 | 15 | 0 | 0 | 17 | 65 | 47 | 21 | 15 | 2 | 2 | 0 | 1 | 23 | 46 | 26 | 0 | 9 | 1 |
| HexNAc(5)(x) | 0 | 0 | 10 | 0 | 1 | 0 | 6 | 0 | 0 | 12 | 0 | 0 | 0 | 7 | 0 | 0 | 0 | 2 | 0 | 0 | 4 | 0 | 0 | 0 |
| HexNAc(5)(F)(x) | 0 | 47 | 2 | 50 | 5 | 52 | 22 | 0 | 0 | 11 | 27 | 28 | 4 | 23 | 2 | 0 | 0 | 0 | 23 | 31 | 10 | 1 | 30 | 1 |
| HexNAc(6+)(x) | 0 | 0 | 1 | 0 | 0 | 0 | 1 | 0 | 0 | 1 | 0 | 0 | 0 | 0 | 0 | 0 | 0 | 0 | 0 | 0 | 2 | 2 | 0 | 0 |
| HexNAc(6+)(F)(x | 0 | 7 | 0 | 10 | 0 | 2 | 0 | 0 | 0 | 0 | 1 | 2 | 0 | 0 | 0 | 0 | 0 | 0 | 5 | 10 | 4 | 91 | 11 | 8 |
| Unoccupied | 100 | 5 | 0 | 18 | 0 | 2 | 1 | 16 | 0 | 0 | 0 | 0 | 0 | 0 | 95 | 0 | 0 | 5 | 17 | 4 | 13 | 5 | 50 | 91 |
| Core | 0 | 4 | 0 | 1 | 0 | 1 | 1 | 5 | 0 | 0 | 0 | 3 | 0 | 0 | 0 | 0 | 0 | 1 | 1 | 0 | 1 | 0 | 0 | 0 |

| | | | | | | | | | | | | | | | | | | | | | | | | |
|---|---|---|---|---|---|---|---|---|---|---|---|---|---|---|---|---|---|---|---|---|---|---|---|---|
| Oligomannose | 0 | 3 | 14 | 1 | 31 | 1 | 38 | 75 | 100 | 5 | 3 | 10 | 43 | 15 | 0 | 98 | 59 | 59 | 18 | 1 | 3 | 0 | 0 | 1 |
| Hybrid | 0 | 0 | 1 | 0 | 25 | 0 | 6 | 2 | 0 | 7 | 1 | 5 | 22 | 21 | 0 | 0 | 38 | 21 | 7 | 1 | 18 | 0 | 0 | 0 |
| Complex | 0 | 91 | 85 | 80 | 44 | 96 | 55 | 7 | 0 | 88 | 96 | 85 | 35 | 64 | 4 | 2 | 3 | 15 | 57 | 94 | 66 | 95 | 50 | 9 |
| Unoccupied | 100 | 5 | 0 | 18 | 0 | 2 | 1 | 16 | 0 | 0 | 0 | 0 | 0 | 0 | 95 | 0 | 0 | 5 | 17 | 4 | 13 | 5 | 50 | 91 |
| Fucose | 0 | 87 | 13 | 79 | 37 | 96 | 40 | 1 | 0 | 29 | 97 | 85 | 35 | 47 | 4 | 2 | 1 | 2 | 58 | 94 | 48 | 93 | 50 | 9 |
| NeuAc | 0 | 28 | 29 | 40 | 20 | 33 | 6 | 0 | 0 | 10 | 32 | 11 | 4 | 18 | 2 | 0 | 8 | 13 | 13 | 17 | 51 | 65 | 50 | 6 |

**Supplemental Table 3: Site-specific glycan analysis of B.1.617.2 (Delta) spike glycoprotein**

| | N61 | N74 | N122 | N149 | N165 | N234 | N282 | N331 | N343 | N603 | N616 | N657 | N709 | N717 | N801 | N1074 | N1098 | N1134 | N1158 | N1173 | N1194 |
|---|---|---|---|---|---|---|---|---|---|---|---|---|---|---|---|---|---|---|---|---|---|
| M9Glc | 0 | 0 | 0 | 0 | 0 | 1 | 0 | 0 | 0 | 0 | 0 | 0 | 0 | 0 | 0 | 0 | 0 | 0 | 0 | 0 | 0 |
| M9 | 0 | 0 | 0 | 0 | 0 | 76 | 0 | 0 | 0 | 0 | 0 | 0 | 0 | 0 | 0 | 0 | 0 | 0 | 0 | 0 | 0 |
| M8 | 0 | 0 | 1 | 0 | 1 | 16 | 0 | 0 | 0 | 1 | 0 | 0 | 0 | 0 | 6 | 0 | 0 | 0 | 0 | 0 | 0 |
| M7 | 0 | 0 | 17 | 0 | 0 | 2 | 0 | 0 | 0 | 8 | 0 | 0 | 100 | 26 | 23 | 0 | 1 | 0 | 0 | 0 | 0 |
| M6 | 0 | 0 | 33 | 0 | 10 | 1 | 0 | 0 | 0 | 13 | 2 | 0 | 0 | 43 | 13 | 0 | 1 | 0 | 0 | 0 | 0 |
| M5 | 28 | 0 | 41 | 0 | 42 | 0 | 3 | 0 | 20 | 62 | 26 | 0 | 0 | 0 | 41 | 9 | 0 | 1 | 0 | 0 | 0 |
| M4 | 4 | 0 | 1 | 0 | 0 | 0 | 0 | 0 | 0 | 4 | 0 | 0 | 0 | 0 | 6 | 0 | 0 | 0 | 0 | 0 | 0 |
| M3 | 1 | 0 | 0 | 0 | 0 | 0 | 0 | 0 | 0 | 0 | 0 | 0 | 0 | 0 | 2 | 0 | 0 | 0 | 0 | 0 | 0 |
| FM | 0 | 0 | 0 | 0 | 0 | 0 | 0 | 0 | 0 | 0 | 0 | 0 | 0 | 0 | 0 | 0 | 0 | 0 | 0 | 0 | 0 |
| Hybrid | 7 | 0 | 5 | 0 | 9 | 0 | 6 | 0 | 0 | 10 | 27 | 0 | 0 | 30 | 7 | 1 | 22 | 3 | 0 | 0 | 0 |
| Fhybrid | 2 | 0 | 0 | 0 | 4 | 0 | 0 | 0 | 11 | 0 | 6 | 0 | 0 | 0 | 0 | 28 | 13 | 4 | 0 | 0 | 0 |
| HexNAc(3)(x) | 4 | 0 | 0 | 0 | 19 | 0 | 4 | 0 | 0 | 1 | 14 | 0 | 0 | 0 | 0 | 0 | 4 | 1 | 0 | 0 | 0 |
| HexNAc(3)(F)(x) | 3 | 0 | 0 | 0 | 0 | 0 | 0 | 0 | 3 | 0 | 2 | 0 | 0 | 0 | 0 | 15 | 3 | 5 | 0 | 0 | 0 |
| HexNAc(4)(x) | 18 | 0 | 0 | 0 | 4 | 0 | 15 | 0 | 0 | 0 | 16 | 0 | 0 | 0 | 0 | 0 | 18 | 0 | 0 | 0 | 0 |
| HexNAc(4)(F)(x) | 7 | 5 | 0 | 8 | 4 | 0 | 7 | 0 | 29 | 0 | 3 | 40 | 0 | 0 | 0 | 25 | 24 | 36 | 0 | 0 | 0 |
| HexNAc(5)(x) | 16 | 0 | 0 | 0 | 3 | 0 | 30 | 0 | 0 | 0 | 6 | 0 | 0 | 0 | 0 | 0 | 5 | 0 | 0 | 0 | 0 |
| HexNAc(5)(F)(x) | 7 | 58 | 0 | 84 | 2 | 0 | 22 | 100 | 36 | 0 | 0 | 46 | 0 | 0 | 0 | 16 | 8 | 34 | 0 | 0 | 0 |
| HexNAc(6+)(x) | 2 | 0 | 0 | 0 | 0 | 0 | 3 | 0 | 0 | 0 | 0 | 0 | 0 | 0 | 0 | 0 | 0 | 0 | 0 | 0 | 0 |
| HexNAc(6+)(F)(x) | 0 | 16 | 0 | 8 | 0 | 0 | 10 | 0 | 0 | 0 | 0 | 5 | 0 | 0 | 0 | 0 | 0 | 14 | 100 | 0 | 15 |
| Unoccupied | 0 | 12 | 0 | 0 | 0 | 3 | 0 | 0 | 0 | 0 | 0 | 7 | 0 | 0 | 0 | 7 | 0 | 2 | 0 | 100 | 84 |
| Core | 1 | 8 | 1 | 0 | 0 | 0 | 0 | 0 | 1 | 0 | 0 | 1 | 0 | 0 | 1 | 0 | 0 | 0 | 0 | 0 | 0 |

| | N61 | N74 | N122 | N149 | N165 | N234 | N282 | N331 | N343 | N603 | N616 | N657 | N709 | N717 | N801 | N1074 | N1098 | N1134 | N1158 | N1173 | N1194 |
|---|---|---|---|---|---|---|---|---|---|---|---|---|---|---|---|---|---|---|---|---|---|
| Oligomannose | 33 | 0 | 93 | 0 | 53 | 96 | 3 | 0 | 20 | 88 | 27 | 0 | 100 | 70 | 90 | 9 | 2 | 1 | 0 | 0 | 0 |
| Hybrid | 10 | 0 | 5 | 0 | 14 | 0 | 6 | 0 | 11 | 10 | 33 | 0 | 0 | 30 | 8 | 29 | 35 | 7 | 0 | 0 | 0 |
| Complex | 56 | 80 | 1 | 100 | 33 | 0 | 91 | 100 | 68 | 1 | 40 | 92 | 0 | 0 | 1 | 55 | 63 | 90 | 100 | 0 | 15 |
| Unoccupied | 0 | 12 | 0 | 0 | 0 | 3 | 0 | 0 | 0 | 0 | 0 | 7 | 0 | 0 | 0 | 7 | 0 | 2 | 0 | 100 | 84 |
| Fucose | 20 | 80 | 1 | 100 | 11 | 0 | 39 | 100 | 78 | 0 | 11 | 92 | 0 | 0 | 0 | 83 | 48 | 93 | 100 | 0 | 15 |
| NeuAc | 9 | 43 | 2 | 97 | 4 | 0 | 17 | 0 | 2 | 0 | 13 | 38 | 0 | 0 | 3 | 7 | 65 | 37 | 74 | 0 | 15 |

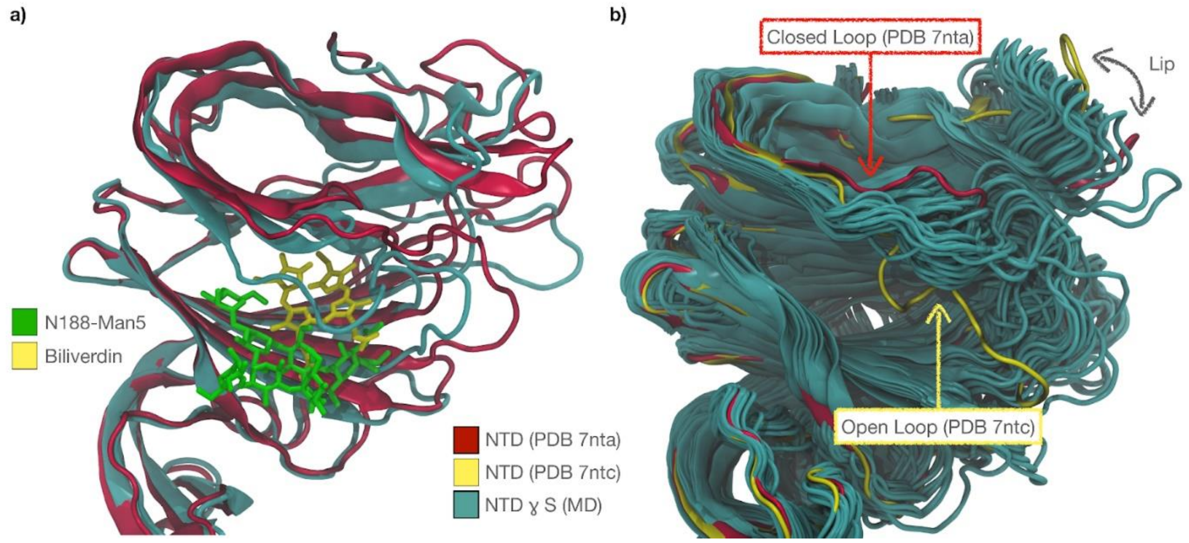


**Supplemental Figure 2. Panel a)** Structural alignment of the NTD from SARS-CoV-2 WH1 S bound to biliverdin (PDB 7nta, red cartoon), and the NTD from the SARS-CoV-2 P.1 S from MD (frame at 793 ns). The backbone root-mean-square deviation (RMSD) value is 1,260 Å for 233 atoms (310 aligned). Biliverdin is represented in yellow sticks and N188-Man5 in green sticks. **Panel b)** Structural alignment of the NTD from SARS-CoV-2 WH1 S bound to biliverdin (PDB 7nta, red cartoon), to the NTD from SARS-CoV-2 Wuhan-hu-1 S bound to the P008_056 Fab (PDB 7ntc, yellow cartoon; antibody not shown), and to the NTD from the SARS-CoV-2 P.1 S from MD, with frames represented every 15 ns steps, from 300 ns to 1.05 µs. Molecular rendering with VMD (https://www.ks.uiuc.edu/Research/vmd/).
